# Fast adaptation of cooperative channels engenders Hopf bifurcations in auditory hair cells

**DOI:** 10.1101/2021.10.11.464021

**Authors:** Francesco Gianoli, Brenna Hogan, Émilien Dilly, Thomas Risler, Andrei S. Kozlov

## Abstract

Since the pioneering work of Thomas Gold published in 1948, it has been known that we owe our sensitive sense of hearing to a process in the inner ear that can amplify incident sounds on a cycle-by-cycle basis. Termed the active process, it uses energy to counteract the viscous dissipation associated with sound-evoked vibrations of the ear’s mechanotransduction apparatus. Despite its importance, the mechanism of the active process and the proximate source of energy that powers it have remained elusive—especially at the high frequencies characteristic of mammalian hearing. This is partly due to our insufficient understanding of the mechanotransduction process in hair cells, the sensory receptors and amplifiers of the inner ear. It has previously been proposed that a cyclical binding of Ca^2+^ ions to individual mechanotransduction channels could power the active process. That model, however, relied on tailored reaction rates that structurally forced the direction of the cycle. Here, we ground our study on our previous model of hair-cell mechanotransduction, which relied on the cooperative gating of pairs of channels, and incorporate into it the cyclical binding of Ca^2+^ ions. With a single binding site per channel and reaction rates drawn from thermodynamic principles, our model shows that hair cells behave as nonlinear oscillators that exhibit Hopf bifurcations, dynamical instabilities long understood to be signatures of the active process. Using realistic parameter values, we find bifurcations at frequencies in the kilohertz range with physiological Ca^2+^ concentrations. In contrast to the myosin-based mechanism, responsible for low-frequency relaxation oscillations in the vestibular hair cells of amphibians, the current model relies on the electrochemical gradient of Ca^2+^ as the only energy source for the active process and on the relative motion of cooperative channels within the stereociliary membrane as the single mechanical driver. Equipped with these two mechanisms, a hair bundle proves capable of operating at frequencies in the kilohertz range, characteristic of mammalian hearing.

**SIGNIFICANCE:** How the inner ear amplifies incident sounds at audible frequencies of several kilohertz is a key question that has remained unanswered despite decades of research into several candidate mechanisms. Here, we model the behavior of hair cells, the sensory receptors of the inner ear, and show that they can undergo oscillatory instabilities called Hopf bifurcations due to the effect of Ca^2+^ on the cooperative opening and closing of mechanotransduction ion channels. Close to the bifurcation point, a hair cell behaves as a nonlinear oscillator that can amplify its input on a cycle-by-cycle basis. We find that our proposed mechanism can operate in the kilohertz range.

## INTRODUCTION

The active process of the inner ear is the vibrating soul of the auditory system (1). The ear employs it to amplify sounds, to sharpen its frequency selectivity, and to help compress six orders of magnitude in sound amplitude into a hundredfold range in the firing rate of the auditory fibers (2). The leading theories trace this process to the biophysical properties of individual hair cells, the sensory receptors of the inner ear. These cells act both as microphones that pick up sound vibrations with their bundles of enlarged microvilli, called stereocilia, and as stimulus amplifiers (3). Both signal transduction and amplification rely on elastic molecular filaments, called tip links, connected to mechanically sensitive ion channels that open and close in unison with hair-bundle vibrations (4). Signal amplification—powered by the active process—requires a source of energy and has to be fast to operate at physiological frequencies, up to 20 kHz in humans and higher in some other mammals.

Although the precise biophysical mechanism at the origin of the active process is still a matter of debate, the ear as a whole—as well as individual hair cells in amphibians—show signatures of dynamical systems operating close to Hopf bifurcation points (5). Such dynamical systems display the most salient features of the active ear, including active amplification of low-intensity stimuli, sharp frequency selectivity, compressive nonlinearity in response amplitude, and spontaneous oscillations (2). Spontaneous oscillations have been proposed to underlie spontaneous otoacoustic emissions—the tones that healthy ears emit when in a quiet environment (6)—and have been measured in hair bundles of the bullfrog sacculus (7). Despite the unifying power of modeling the ear as a collection of nonlinear oscillators close to Hopf bifurcation points, the connection between the cellular and molecular biology of the inner ear and this overarching mathematical description has not been fully understood.

Current models of mechanotransduction reproduce well the behavior of the hair bundles of the bullfrog sacculus, which exhibit all features of the active process as they detect and amplify stimuli at frequencies on the order of tens to a hundred hertz (5, 8). These models, however, struggle to explain how auditory hair cells can achieve active amplification at frequencies in the kilohertz range, typical of mammalian hearing. This stems from the fact that such models are limited by the pace of myosin motors on which they rely to reset the working range of the hair cell on a cycle-by-cycle basis (5). Despite several pioneering studies on this subject (8–11), no microscopic model of hair-bundle motility has so far been able to reproduce the properties of active amplification at frequencies in the kilohertz range with only a minimal number of known, possible states of the channels, as well as realistic values of the gating swing, the amplitude of the conformational change of an individual channel upon opening or closing. In mammals, sparse evidence suggests that hair bundles are capable of generating forces (12) that could drive the active process; yet, no mammalian hair cell has ever been observed to oscillate spontaneously. The source of amplification in the cochlea has rather been attributed to electromotiliy, the ability of mammalian outer hair cells to change their length in response to changes in their transmembrane potential (13). Unlike the low-frequency hair-bundle motility observed in hair cells of the bullfrog sacculus, electromotility would in principle be fast enough to drive the active process at higher frequencies. However, although essential for mammalian hearing, electromotility alone cannot account for the active process because it is mostly linear at small displacements, and because the hallmarks of the active process persist under conditions where electromotility vanishes (14–16).

Here, we present a new model of hair-bundle motility that relies on the established electrochemical gradient of Ca^2+^ as an energy source to power active oscillations, similarly to what has been proposed by Choe *et al.,* (1998) (9). We base our model on an earlier proposal that each tip link is connected to two channels, the states of which are reciprocally coupled to hair-bundle motion by membrane-mediated forces (17, 18). This model is not limited by the action of myosin motors and can show spontaneous oscillations at high frequencies. With two channels per tip link, which can move along the lipid bilayer, the model relies on only one Ca^2+^ binding site per channel, on a realistic value for the individual-channel gating swing, and on transition rates derived from thermodynamic principles. We describe the details of this model in the next section.

## MODEL

Based on previous work (17), we hypothesize that in a mature hair bundle every tip link is connected to two functional mechanosensitive ion channels (19, 20). These channels are mobile in the membrane and they each connect to one of the two strands of the tip link at its lower end (21). Two adaptation springs anchor the channels to the cytoskeleton (see Fig. 1 and Fig. 2A). Although initially a conjecture (17), this connection has later been confirmed experimentally (22). The two channels function as a unit due to a reciprocal coupling between their conformational states (open or closed) and the shape of the membrane in which they sit (23–25). Minimizing the bending energy of the membrane surrounding the channels results in cooperative gating, such that only two gating states (open-open and closed-closed) are effectively at play for the pair. In contrast to the classical gating-spring model (4, 26), this configuration does not require a large gating swing to explain the observed relaxation of the hair bundle upon channel opening. In this two-channel model, the relaxation is instead provided by the motion of the channels along the membrane.

**Figure 1:**
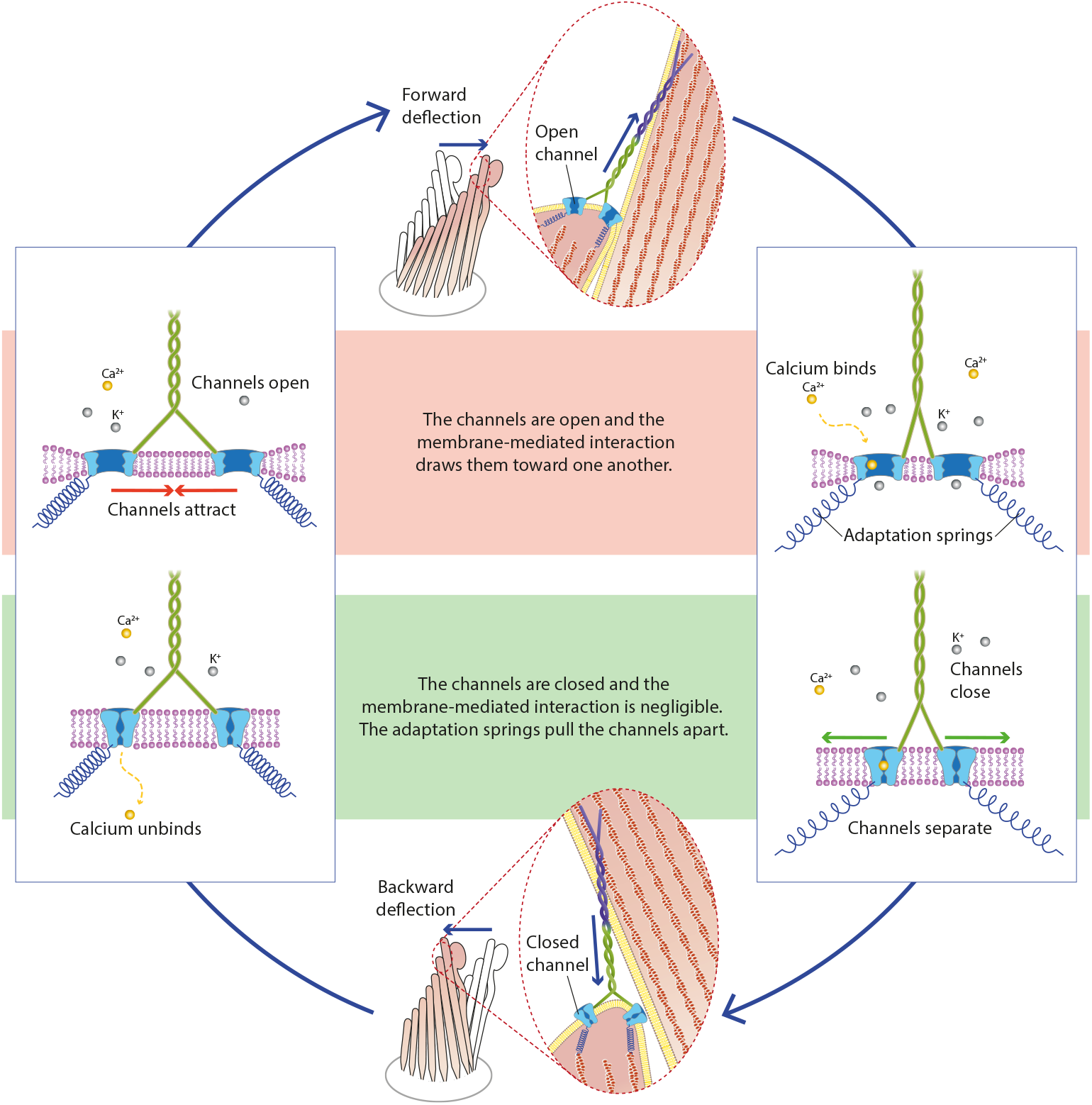
Schematics of the hair-bundle oscillations driven by Ca^2+^ binding and unbinding (bundle deflections are exaggerated for the purpose of the illustration). We start the description of the cycle from the left box, upper drawing, with both channels open and no Ca^2+^ bound. In that configuration, the membrane-mediated interaction between the two channels drives them toward one another (left box, upper panel, red arrows). This movement relaxes tension in the tip link, therefore allowing the stereociliary pivots to drive the hair bundle to the right (top illustration, blue arrows). The channels being open, Ca^2+^ enters and subsequently binds to one of the two channels, facilitating its closure. Since the open-closed state is strongly disfavored energetically due to the membrane-mediated interaction between the channels, this triggers the closure of the second channel almost simultaneously (right box, lower panel). In this closed-closed configuration, the membrane potential is almost flat and tension in the adaptation springs pulls the channels apart (green arrows). The resulting increase of tension in the tip link causes the bundle to twitch backwards (bottom illustration, blue arrows). Now that the channels are closed, Ca^2+^ diffuses within the cell and its local concentration decreases, which favors Ca^2+^ unbinding (left box, lower panel). Tension in the springs then tends to reopen the channels, thus bringing us where we started and completing the cycle of oscillations.

Here, we add to that description an active process that is based on a cycle of Ca^2+^ binding and unbinding to the channels, similarly to what was proposed by Choe *et al.* (9) in 1998. We hypothesize that the binding of one Ca^2+^ ion to a channel lowers the energy of the closed state, while leaving the open state unaffected. We further hypothesize that the rate of Ca^2+^ binding depends on its local concentration, whereas Ca^2+^ unbinding occurs at a constant rate. Since Ca^2+^ local diffusion occurs on much shorter timescales than those of interest here, we estimate that the Ca^2+^ concentration at the channel binding site equilibrates instantaneously to two values, one for each channel state (open or closed). We therefore introduce two rates of Ca^2+^ binding, thereby breaking thermodynamic equilibrium. Such an active process is shown to be able to drive spontaneous oscillations of the hair bundle (Fig. 1). In response to a change of parameter values, and notably on Ca^2+^ concentration in the vicinity of the open channel, the hair bundle transitions between a quiescent and an oscillatory dynamical regime. This transition corresponds to a Hopf bifurcation, where the system’s sensitivity is maximal.

### Force balance

This part of the model is identical to the model published in (17). Force balance on the hair bundle reads:

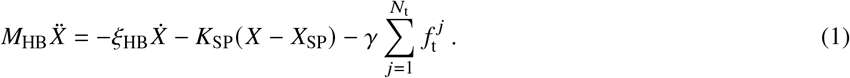

Here, *X* denotes the position of the tip of the hair bundle, projected onto the direction of mechanosensitivity, 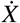 and 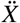 are its first and second time derivatives, and *X*_SP_ is the resting position of the hair bundle in the absence of tip links, due to the action of the stereociliary pivots; *M*_HB_ is the hair bundle’s apparent wet mass (9, 27), *ξ*_HB_ the bundle’s viscous drag coefficient, *K*_SP_ the combined stiffness of the stereociliary pivots, and *N*_t_ the number of tip links. The scaling factor *γ* is the approximately constant projection factor of the tip-link axis on the hair-bundle displacement axis, such that for a change of position of the hair bundle *X*, the tip-link extension changes by *x = γX*. Each of the *N*_t_ tip links in the hair bundle contributes a projected force 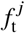 along the tip-link axis given by

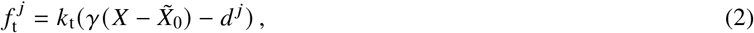

where *k*_t_ is the tip-link stiffness and *d^j^* the varying distance between the tip-link fork and the membrane of the lower stereocilium along the tip-link axis. An illustration of the corresponding geometry is shown in Fig. 2A. The displacement 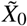 is a reference position of the *X*-axis that sets a reference tension in the tip links for a given hair-bundle position *X*. This tension stems from the action of myosin motors, which incessantly pull on the tip link’s upper insertion point (28). Focusing on shorter timescales than those associated with myosin-motor displacements, we consider 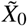 as a constant quantity in the following.

**Figure 2.**
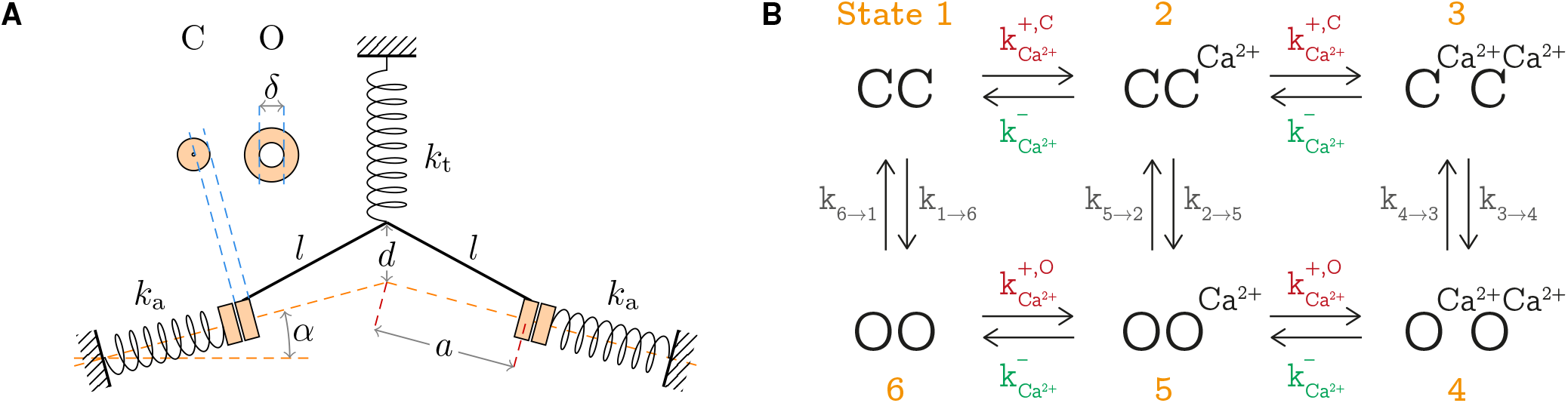
(A) Geometry of the two-channel model and associated parameters. The orange doughnuts represent the mechanosensitive channel in the closed (C) and open (O) configuration. The system of springs is shown in the CC configuration. (B) Schematic representation of the six different states considered for a channel pair. States 1, 2, and 3 correspond to both channels closed, with respectively either zero, one, or two calcium ions bound. States 4, 5, and 6 correspond to both channels open, similarly with different numbers of calcium ions. Each allowed direct transition is characterized by rate constant, where the transitions 1 ⇌ 6, 2 ⇌ 5, and 3 ⇌ 4 correspond to opening or closing of the channel, and 1 ⇌ 2, 2 ⇌ 3, 4 ⇌ 5, and 5 ⇌ 6 correspond to binding or unbinding of Ca^2+^.

To close the system of equations, we now need to determine how the distance *d^j^* relates to the hair-bundle displacement *X*, for each tip link *j*. Force balance on either of the two channels connected to the tip link reads:

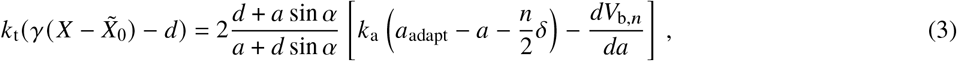

where we define *a* as half the distance between the two tip-link insertion points into their respective channels, projected on the membrane plane, *α* is the angle between the perpendicular to the tip-link axis and the stereociliary-membrane plane, *k*_a_ is the adaptation springs’ stiffness, **a**_adapt_ sets the inter-channel distance for which the adaptation springs slacken, *δ* is the single-channel gating swing, *n* = 0 1, or 2 is the number of opened channels in the pair, and *V*_b,*n*_ is the elastic potential of the stereociliary lipid bilayer in which the mechanosensitive channels are embedded, which depends on *n* (for an illustration see Fig. 2A). The mathematical expression of this elastic membrane potential, together with associated parameters, is given in ref. (17). It is not explicitly reported here for the sake of conciseness. Finally, geometrical constraints impose that *d* and *a* are related by:

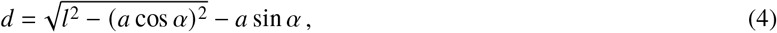

where *l* is the length of the tip-link fork. A full description of that part of the model together with detailed graphical illustrations are given in (17), and parameter values used in the present study are provided in Table 1.

**Table 1:** Model’s parameters.

|  | Parameter | Description | Value | Source |
| --- | --- | --- | --- | --- |
| Mechanics | $\alpha$ | Angle of adaptation springs with respect to horizontal | $15^\circ$ | |
| | $K_{SP,0}$ | Reference combined stiffness of the stereociliary pivots | $0.65 \text{ mN} \cdot \text{m}^{-1}$ | Jaramillo and Hudspeth (30) |
| | $\gamma_0$ | Reference geometrical scaling factor | 0.12 | Choe et al. (9) |
| | $N_t$ | Number of tip links | $7/8 \cdot N_s$ | Choe et al. (9) |
| | $M_{HB}$ | 'Wet mass' of the hair bundle | 100 pg | Baumgart (27) |
| | $\xi_{HB}$ | Friction coefficient of the hair bundle | $100 \text{ nN} \cdot \text{s} \cdot \text{m}^{-1}$ | Choe et al. (9) |
| Calcium cycle | $E_{Ca^{2+}}$ | Energy associated to $Ca^{2+}$ binding/unbinding (closed state) | $1 k_B T$ | adapted from Beurg et al. (31) |
| | $k_b$ | Reaction constant of $Ca^{2+}$ binding/unbinding | $1 \text{ ms}^{-1} \cdot \mu\text{M}^{-1}$ | Cheung and Corey (32) |
| | $\bar{k}$ | Attempt frequency for channel opening and closing | $10^4 \text{ s}^{-1}$ | Beurg et al. (31) |
| | $K_d$ | Reference $Ca^{2+}$ concentration for $Ca^{2+}$ unbinding | $20 \mu\text{M}$ | Cheung and Corey (32) |
| | $[Ca^{2+}]^{\text{close}}$ | $Ca^{2+}$ concentration at the binding site of a closed channel | $0.05 \mu\text{M}$ | Choe et al. (9) |
| | $[Ca^{2+}]^{\text{open}}$ | $Ca^{2+}$ concentration at the binding site of an open channel | $37 \mu\text{M}$ | Choe et al. (9) |

### Calcium cycle

We describe here the implementation of the influence of Ca^2+^ binding and unbinding to the channels on their gating states. We hypothesize one Ca^2+^ binding site per channel, which leads to a total of three Ca^2+^-binding states for a given pair of channels, with a total of either zero, one, or two ions bound to the channel pair. The rates of Ca^2+^ binding depend on the local Ca^2+^ concentration. We consider that this concentration is set by the channel’s state, open or closed, since it has been shown that this concentration reaches its steady-state value on sub-millisecond timescales after channel opening or closing (9, 29). The rate of Ca^2+^ unbinding, however, is supposed to be independent of the local concentration. This leads to:

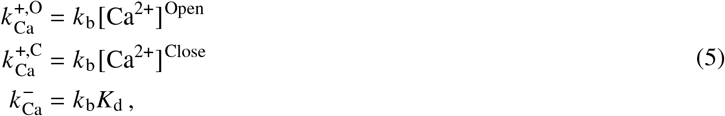

where 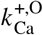 and 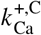 are the rates of Ca^2+^ binding in the open and closed states, respectively, 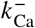 is the rate of Ca^2+^ unbinding, *k*_b_ is the reaction constant of Ca^2+^ binding, and *K*_d_ is a concentration that sets the rate of Ca^2+^ unbinding.

### Opening and closing transition rates

To complete the model, we now focus on identifying the probabilities of the different states that a channel pair can occupy, and how they evolve with time. With physiological parameters, due to the divergence of the membrane elastic potential at small inter-channel distances, the energies of the OC states are higher than those of either OO or CC states by several *k*_B_*T*. In the following, we therefore ignore the occupancy of this state and investigate only the transitions between the OO and CC states. This corresponds to investigating a perfect cooperativity between the two channels, mediated by the lipid bilayer (17). Taking into account the different occupancy states of the channels by Ca^2+^, we are therefore left with a total of six states, with three Ca^2+^ binding states for each of the two gating states of the channel pair.

Transitions between each of these adjacent states in this calcium cycle occur with specific rates, schematically represented in Fig. 2B. Among these rates, the Ca^2+^ binding and unbinding rates have already been discussed, and we need to specify the transition rates between the OO and CC states with a given number of bound Ca^2+^ ions. To write these rates, we suppose that each of these transitions goes through its respective OC state, which serves as a limiting transition state. The associated activation energy allows to write the transition rate using Kramers’ reaction-rate theory. We hypothesize that the presence of one calcium ion bound to a closed channel lowers its energy by the fixed amount *E*_Ca_. The energies of each of the six states represented in Fig. 2 therefore read:

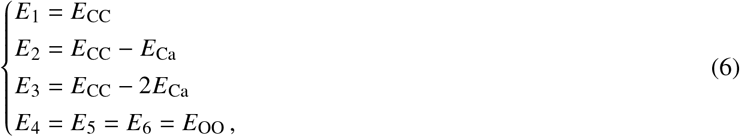

where *E*_CC_ and *E*_OO_ are respectively the energies of the CC and OO states in the absence of calcium ions. The energies *E*_1⇌6_, *E*_2⇌5_, and *E*_3⇌4_ of the respective transition states between the states 1 and 6, 2 and 5, and 3 and 4, in both directions, read:

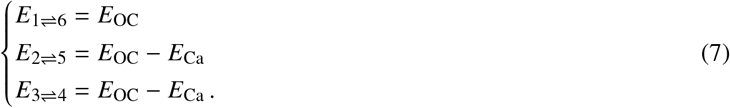

For the transition state between states 2 and 5, there are *a priori* two possibilities: OCa–C and O–CCa. As we know that the closed state is stabilized by calcium, the intermediate state O–CCa is of lower energy. Due to the exponential dependence of the transition rate on the transition-state energy, we consider that the state O–CCa is strongly favored, leading to the expression given above. Note that we assume that the binding of Ca^2+^ does not affect the energy of the open state. As a result, the height of the energy barrier in the transitions 5 → 2 and 4 → 3 is lowered by *E*_Ca_ in both cases as compared with that of the transition 6 → 1. The energy barriers in the opposite directions, however, behave differently: the barrier of the transition 2 → 5 is unchanged as compared to that of 1 → 6, and that of 3 → 4 is increased by *E*_Ca_. Given these energies, Kramers’ rate theory allows us to write the following expressions for the forward and reverse transition rates between the states 1 and 6, 2 and 5, and 3 and 4:

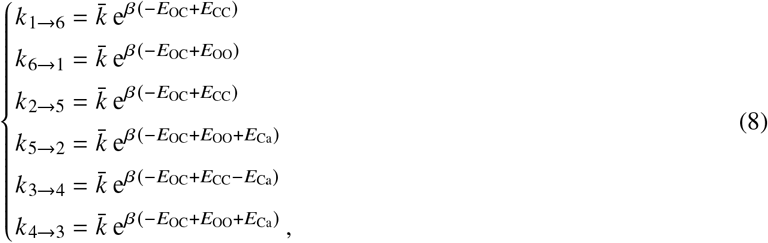

where 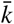 is a pre-exponential factor called ‘the attempt frequency’ that sets a baseline rate to these transitions.

### Probability dynamics

We can now write the time-evolution equations for the occupancy probabilities of each of the six different states shown in Fig. 2:

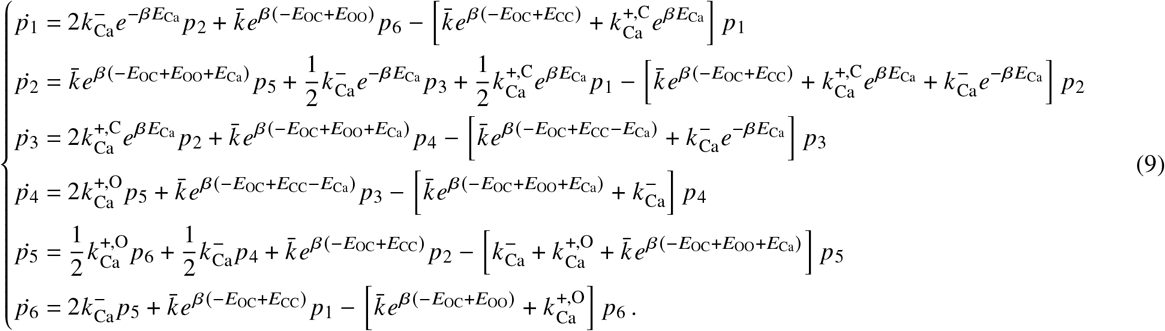

Note that the prefactors 2 and 1/2 that appear in some terms are due to the fact that states 2 and 5 are degenerate: in both instances, a single Ca^2+^ ion is free to bind to either of the two available channels, with the same resultant energy in either case.

### Expression of the energies

In the system of equations 9, the energies *E*_OO_, *E*_OC_, and *E*_CC_ correspond to the energies of one pair of channels in the respective configurations. These energies include the elastic contributions from the extensions of the tip link and adaptation-springs, that of the lipid-bilayer deformations *V*_b,*n*_, as well as the gating energy of the channels *E*_g_, the energy difference between the open and closed states of a single channel. We define the following energy function for a pair of *n* = 0, 1, or 2 open channels, located a distance 2*a* apart, the hair bundle being positioned at *X*:

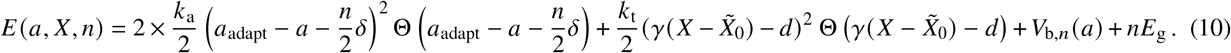

Here, *d* is a function of the half inter-channel distance *a* thanks to the geometric relation 4, and **Θ** is the Heaviside step function that represents slacking, since neither the tip links nor the adaptation springs are supposed to resist compression. The energies *E*_OO_, *E*_OC_, and *E*_CC_ in Eq. 9 equal *E*(*a, X, n*) as given in Eq. 10 with *n* = 0, 1, or 2, respectively.

As seen in Eq. 3, together with Eq. 4, the position of the hair bundle’s tip *X* is a function of the half inter-channel distance *a* as well as of the number *n* of opened channels in the pair via the elastic membrane potential *V*_b,*n*_ We therefore have three different relations *X*(*a*)—one per gating state of the pair (OO, OC, or CC). Each of these relations is invertible, such that we can write the equations in terms of the sole variable *X*, replacing the variable *a* in state *n* in Eq. 10 by the corresponding solution to Eq. 3, *a_n_* (*X*). Finally, we obtain expressions that can be summarized under the form:

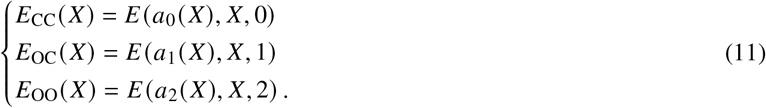

### Closing the system

Considering that Eq. 1 is second order, we can split it into two first-order equations by introducing the variable 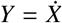. Doing so, we use the vector **S** = (*X, Y, p*_1_, *p*_2_, *p*_3_, *p*_4_, *p*_5_, *p*_6_)^*T*^ to represent the state of the system at any given time, where the symbol ‘T’ corresponds to the matrix transposition operation to get a vector. The whole model now reads:

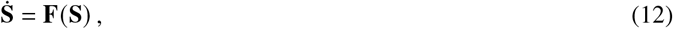

where the vectorial function **F** corresponds to Eqs. 1 and 9, together with the other equations given above that allow to express **F** as a function of the components of **S** only.

### Parameter values

Several of the model parameters can be estimated from the literature. We list in Table 1 the parameters that relate to the mechanics of the hair bundle and to the calcium cycle. The parameters that do not appear in Table 1 are the same as the default values in ref. (17). The parameter 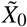 entering Eq. 2 is however not directly observable from experiments, as it is linked to the position of myosin motors within stereocilia. To infer its value, we first choose that the origin of the *X* axis (*X* = 0) is such that it is a resting point of the hair bundle, i.e., a static solution of Eq. 1. This fixes the value of *X*_SP_, the resting position of the stereociliary pivots relative to the origin, provided that we know all the other parameters in that equation. To determine the value of 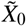, we impose in addition that, at *X* = 0, the steady-state open probability of the channels *P*_open_ = *p*_4_ + 2*p*_5_ + *p*_6_ equals 0.15. This value corresponds to the resting open probability of mammalian inner hair cells tuned to high frequencies (33–36).

## RESULTS

Having all parameters at hand, we perform in the following linear-stability analysis around the fixed point *X* = 0 on the dynamics given by Eq. 12. In the following figures, we plot the real and imaginary components of the pair of eigenvalues of maximum real part, which characterize the dynamics of our system in the proximity of the steady state. As this pair of complex-conjugated eigenvalues crosses the imaginary axis, an oscillatory decay becomes a limit-cycle oscillation. At the precise parameter values for which the real part changes sign, there is a Hopf bifurcation point (37). All the other eigenvalues have negative real parts. They correspond to modes that relax on a finite timescale at the bifurcation point. More generally, looking at the long-time dynamics of the hair bundle, only the pair of eigenvalues of largest real component controls its behavior.

We start by investigating the dependence of the eigenvalues on the number of stereocilia *N*_s_ in the hair bundle. Along the tonotopic axis of the cochlea, hair cells tuned to higher frequencies have more stereocilia than those tuned to lower frequencies, and their stereocilia tend to be shorter, which renders their hair bundles stiffer (38–40). In addition, the total mass of actin contained in each hair bundle of a given cochlea has been shown to be roughly constant along the chicken’s cochlea (41). This suggests a relationship between the geometric gain factor *γ*, the collective stereociliary pivot stiffness *K*_SP_, and the number of stereocilia in the bundle *N*_s_ (9). Specifically, we follow ref (9) and assume that *γ* ∝ *N*_s_ and 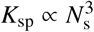, therefore reducing the number of free parameters in the model. We postulate therefore that *K*_SP_ = *K*_SP,0_ · (*N*_s_/50)^3^ and *γ* = *γ*_0_ · (*N*_s_/50), where *K*_SP,0_ and *γ*_0_ are a reference stiffness of the stereociliary pivots and a reference scaling factor in a hair bundle with 50 stereocilia. Their values are set to correspond to those used in (17).

We investigate the dependence of the characteristic frequency of the modeled hair bundle on *N*_s_, following the constant actin-mass hypothesis described above, to compare our results with those of ref. (9). We plot in Fig. 3 this dependence for three values of the attempt frequency 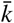. For small values of 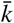, the hair bundle remains quiescent, regardless of the number of stereocilia *N*_s_ (Fig. 3A). For large values of 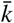, the hair bundle oscillates above a critical value of *N*_s_ (Fig. 3 C). At intermediate values of 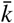, the hair bundle oscillates within a limited range of *N*_s_ values (Fig. 3 B). The characteristic frequency of spontaneous oscillations increases with the number of stereocilia, as in ref. (9).

**Figure 3:**
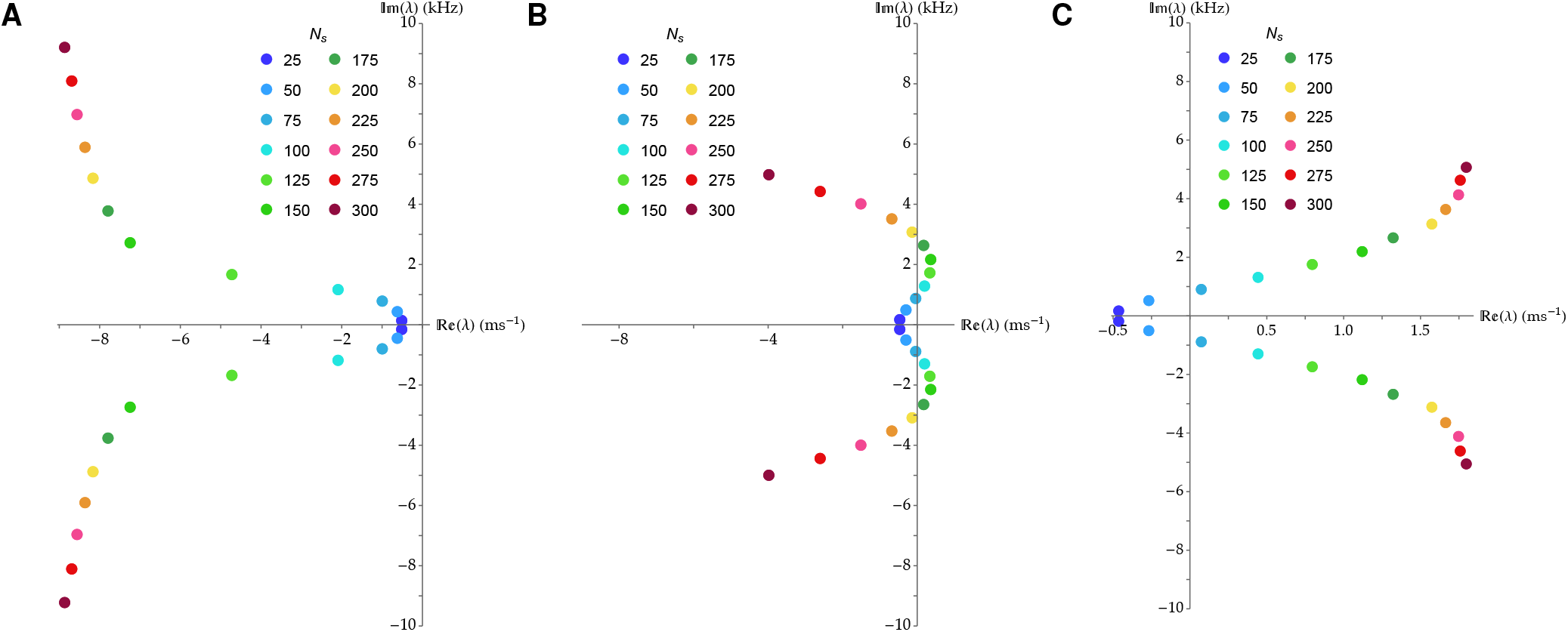
Dependence of the pair of eigenvalues *λ* with the largest real part on the number of stereocilia *N*_s_, under the constant actin-mass hypothesis and for three values of the attempt frequency 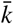. (A) For 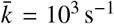, the hair bundle displays only damped oscillations. (B) For 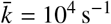, the hair bundle shows limit-cycle oscillations for a limited range of values of *N*_s_, here roughly between 50 and 200. (C) For 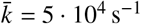, the hair bundle presents limit-cycle oscillations above a threshold value of *N*_s_, here around 50.

### Ca^2+^ dependency

Hair-bundle oscillations are known to be affected by the concentration of calcium in the surrounding milieu (7). In Fig. 4, we show the dependence of the eigenvalues on calcium concentration at the binding site when the channels are open, that is varying the parameter [Ca^2+^]^open^, as well as on the open probability of the channels at steady state *P*_open_. We display in Fig. 4A the dependence of the eigenvalues as a function of the number of stereocilia *N*_s_, for four values of the calcium concentration in the vicinity of the channel in the open state [Ca^2+^]^open^. As argued by Choe *et al.* (9), we can expect the Ca^2+^ concentration close to the channel’s pore to rapidly equilibrate with that of the endolymph upon channel opening—on timescales of the order of a few microseconds—and to return to concentrations a hundred-fold smaller within a few tens of microseconds. Based on these estimates, we can conclude that our parameter [Ca^2+^]^open^ corresponds to the endolymph Ca^2+^ concentration up to frequencies of tens of kilohertz.

We see that the system transitions from damped to limit-cycle and then back to damped oscillations as [Ca^2+^]^open^ increases. The system displays limit-cycle oscillations for calcium concentrations around 40 *μ*M. For lower and higher values of [Ca^2+^]^open^, the eigenvalues are confined to the left of the imaginary axis, in the regime of damped oscillations. The evolution of the eigenvalues with [Ca^2+^]^open^ is shown in Fig. 4B at fixed *N*_s_ = 100. We can see that the variation in characteristic frequency remains relatively small.

As the impact of calcium binding on hair-bundle oscillations depends, at any given time, on the total number of open channels, we expect the behavior of the hair bundle to change as we vary the value of the open probability at steady state. In Fig. 4C, we show the eigenvalues’ dependence on *N*_s_ similarly to Fig. 4A, for six different values of *P*_open_. With the parameter values given in Table 1, the hair bundle displays limit-cycle oscillations within a limited range of *P*_open_ values, from about 0.15 to about 0.30.

**Figure 4:**
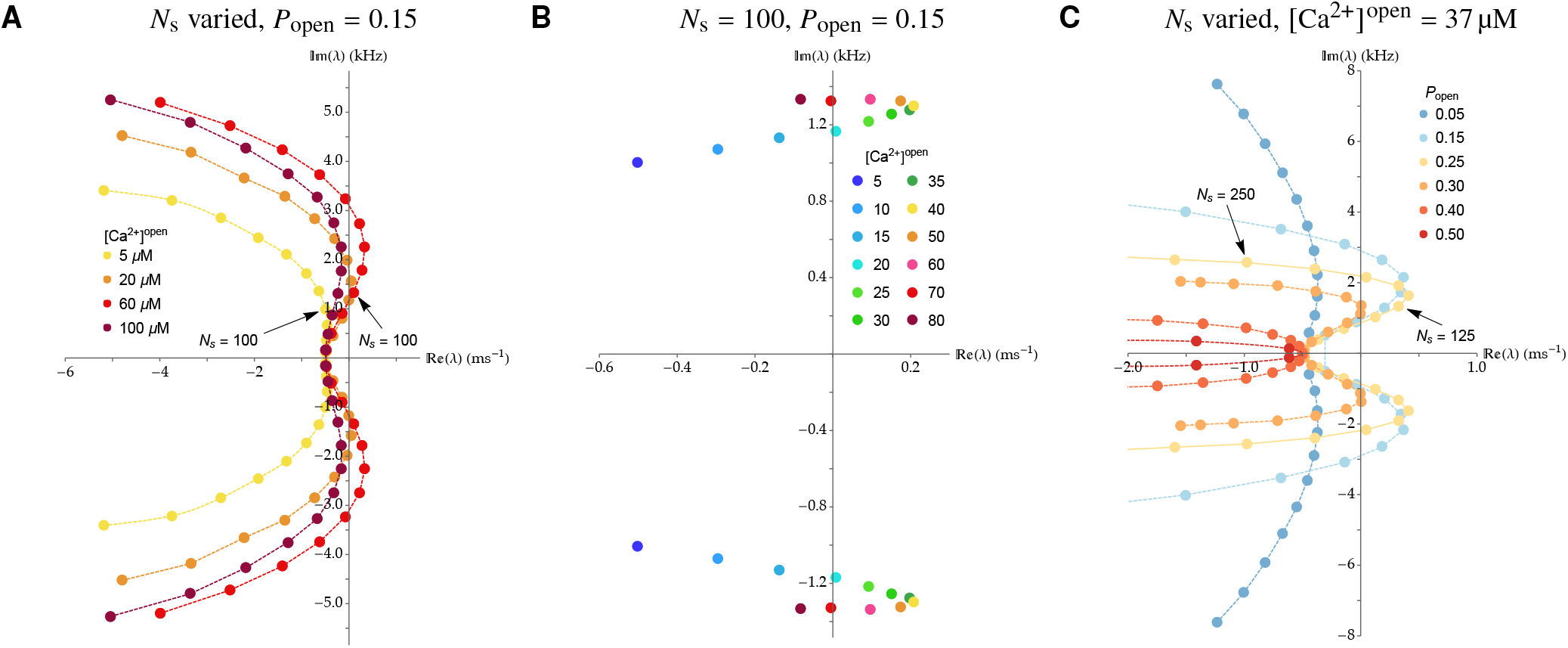
Dependence of the most unstable pair of eigenvalues *λ* on the number of stereocilia *N*_s_, the calcium concentration in the vicinity of the channel in the open state [Ca^2+^]^open^, and the open probability at rest *P*_open_. (A) The eigenvalues are plotted in four families of curves, each corresponding to a different value of [Ca^2+^]^open^, while varying the number of stereocilia *N*_s_ from 25 to 300 as in Fig 3 and at fixed 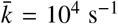 and *P*_open_ = 0.15. (B) The eigenvalues are plotted for various calcium concentrations [Ca^2+^]^open^ at fixed *N*_s_ = 100, 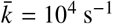, and *P*_open_ = 0.15. (C) The eigenvalues are plotted similarly to panel A, in six families of curves, each corresponding to a different value of *P*_open_, for [Ca^2+^]^open^ = 37 μM, 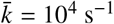, and with the same values of *N*_s_ as in panel A.

In Fig. 5, we represent the frequencies of the spontaneous oscillations, when they exist, in three dimensions as a function of the parameters *K*_SP,0_, *N*_s_, and *k*_t_. Each bubble corresponds to an oscillatory state associated with a unique triplet of these parameters. Both its color and size code for the frequency of the corresponding spontaneous oscillations.

**Figure 5:**
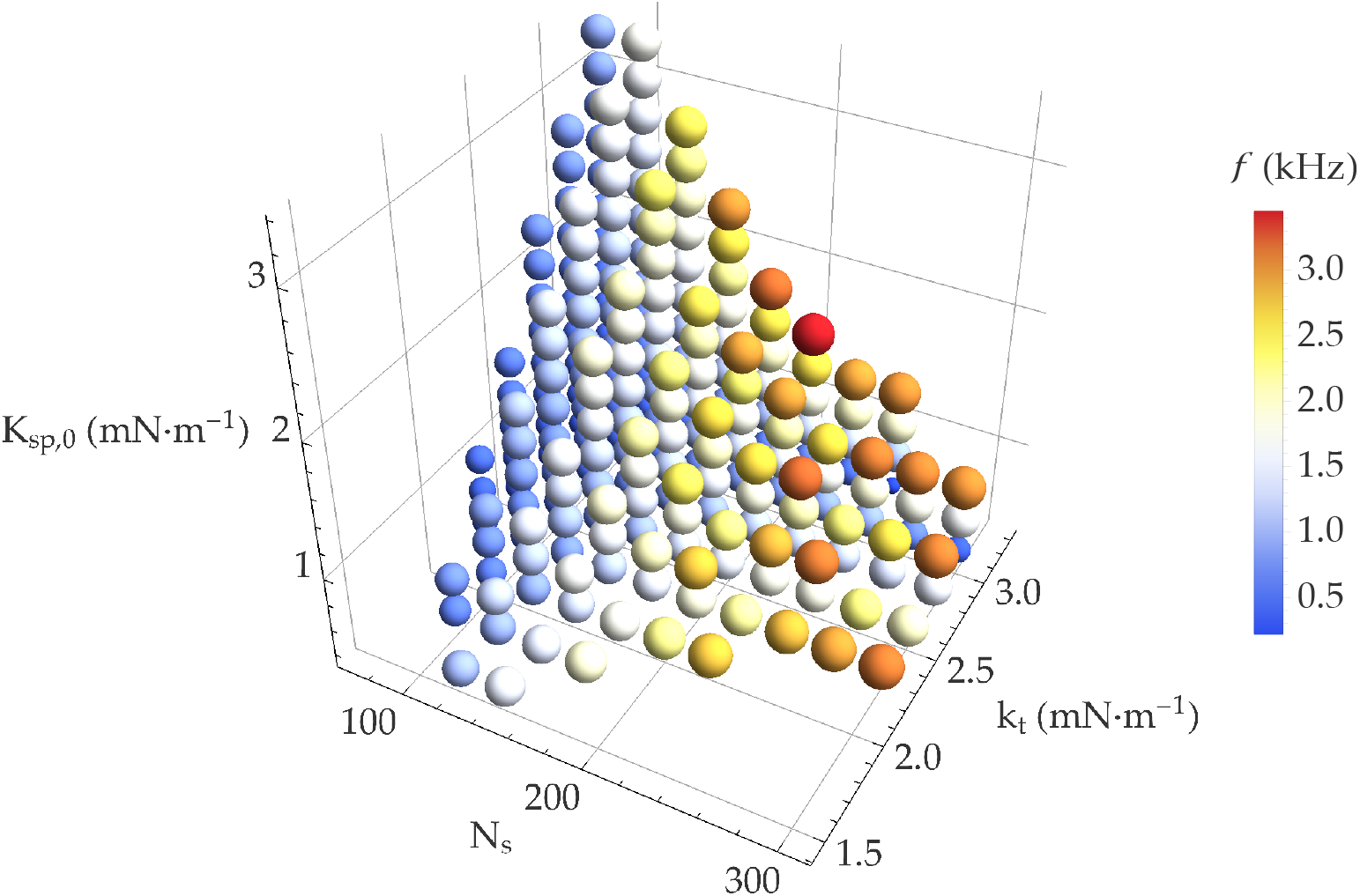
4D plot of the parameter values for which the system shows limit-cycle oscillations as a function of *k*_t_, *N*_s_ and *K*_SP,0_, with 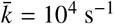 and *P*_open_ = 0.15. Each plotted bubble corresponds to an oscillatory state. As the corresponding frequency varies from 0.2 kHz to 3.5 kHz, the color of the bubbles changes from blue to red and their size increases.

## DISCUSSION

In our previous work, we developed a new gating-spring model with cooperative channels (17) and then used it to explain changes observed in the biophysical properties of hair cells during development and tip-link regeneration (18). In the present paper, we have developed the model further and shown how the mechanotransduction apparatus of a hair cell can not only transduce, but also amplify incident sounds at high frequencies. Specifically, we investigated whether, within this framework, the electrochemical gradient of Ca^2+^ can power spontaneous oscillations in hair cells at high frequencies.

The proposal that the electrochemical gradient of Ca^2+^ ions can power the active process through their binding and unbinding to the mechanosensitive channels is not new. Our work was particularly influenced by that of Choe *et al.* (9), which showed that a Hopf bifurcation can arise from the Ca^2+^-dependent fast adaptation mechanism, without the need for myosin-motor activity. There are, however, major differences between the two models. In the work by Choe *et al.,* all the ‘forward’ rate constants were identical, as were their symmetric ‘reverse’ rate constants. Taking different values for these two rate constants therefore drove the cycle of hair-bundle oscillations. Here, instead, the rate constants are based on Kramers’ rate theory, using the open-closed state of the channel pair as the activation state. The oscillations are driven by the electrochemical gradient of Ca^2+^, as described by the out-of-equilibrium rates of Ca^2+^ binding and unbinding in Eq. 5. The second main difference is that here a single calcium binding site per channel suffices to give rise to spontaneous oscillations, whereas two binding sites were required in the work by Choe *et al.* for energetic and kinetic considerations. In our model, in contrast, the rate constants are set differently, as explained above. Finally, due to the mobility of the channels within the lipid membrane, our model generates spontaneous oscillations with a smaller, more realistic, single-channel gating swing of 2 nm, whereas 4.5 nm was used in the study by Choe *et al..*

Our results are consistent with a number of experimental findings: First, as shown in Fig. 5, the frequency of spontaneous oscillations increases with the number of stereocilia, in agreement with the tonotopic arrangement along the cochlea (40, 41). Second, Tobin *et al.* showed that all three parameters, *N*_s_, *k*_t_ and *K*_sp_ increase along the tonotopic axis of the cochlea (40). This is in agreement with the results shown in Fig. 5, where the characteristic frequency of oscillations increases as these parameters are increased simultaneously. Third, spontaneous oscillations in the model arise only in a limited range of Ca^2+^ concentration. With physiological parameters, this range is approximately 20–70 *μ*M, in agreement with the Ca^2+^ concentration of the cochlear endolymph (42).

With the parameters used in the present study, we have observed the regime of limit-cycle oscillations within a range of open probability at steady state of 10% to 35%. Inner hair cells tuned to high frequencies maintain their resting open probability in the range of 10% to 20% at physiological Ca^2+^ concentration (35, 36). Due to the shape of the sigmoidal open-probability vs. displacement curve, a given high-frequency sinusoidal stimulus results in a greater integral voltage response at such smaller open probabilities (35). We conclude from these results that the range of resting open probabilities characteristic of inner hair cells not only maximizes the integrated voltage response for a given high-frequency input but is also compatible with mechanical amplification.

In the experimentally observed and extensively modeled low-frequency oscillations of hair cells from the bullfrog sacculus, the myosin motors play a direct role in the oscillations. The hair-bundle negative stiffness and the motor-based slow adaptation together engender large-amplitude oscillations in those vestibular hair cells tuned to frequencies of several hertz to tens of hertz. During a complete cycle of these relaxation oscillations, the channels’ open probability varies extensively, from almost zero to about one. In contrast, myosin motors do not play a direct role in the amplification mechanism we are proposing. Here, their role is implicitly limited to setting the channels’ open probability at steady state to a specific value, which they are known to do by means of a process called ‘slow adaptation’. Our results indicate that large-amplitude oscillations are neither expected from the proposed amplification mechanism nor required to support the active process in general. The hair-bundle stiffness does not need to be negative, and the open probability can remain near its target value at all times. This is because sufficiently close to the instability threshold of a supercritical Hopf bifurcation, the oscillation amplitude is arbitrarily small. This could be the reason why, experimentally, one has never observed spontaneous oscillations in cochlear hair cells tuned to frequencies in the kilohertz range. It is possible that spontaneous oscillations in these cells remain so small that they are practically unobservable due to Brownian motion and measurement noise. Therefore, despite the absence of any direct experimental evidence so far, it is possible that auditory hair cells act as cochlear amplifiers at kilohertz frequencies through the mechanism described in this study.

## AUTHOR CONTRIBUTIONS

A.K. conceived and designed the project; T.R. and A.K. directed the project; E.D. and T.R. designed the model; F.G., B.H., E.D., and T.R. wrote the code; F.G. produced the results; F.G., T.R., and A.K. interpreted the results and wrote the paper.

## ACKNOWLEDGMENTS

Work on this project in the Kozlov lab was funded by the Wellcome Trust (108034/Z/15/Z and 214234/Z/18/Z) and the Imperial College Network of Excellence Award.

